# Normalization of aperiodic ECoG components indicates fine motor recovery after sensory cortical stroke in mice

**DOI:** 10.1101/2022.02.16.480472

**Authors:** Jonatan Biskamp, Sara Isla Cainzos, Focko L. Higgen, Christian Gerloff, Tim Magnus

## Abstract

Electrophysiological signatures of ischemic stroke might help to develop a deeper understanding of the mechanisms of recovery. Here, analyses of multichannel electrocorticography (ECoG) in awake mice demonstrated that the shape of power spectral density (PSD) is modulated in the vicinity of sensory cortical stroke. PSD consists of both rhythmic oscillatory and non-rhythmic, aperiodic components. The alteration of spectrum shape was reflected in a transient increase of aperiodic exponents, while the relative power and frequency of slow oscillations remained unchanged in the peri-infarct cortex. Exponents derived from motor areas significantly correlated with recovery of fine motor deficits of the contralateral forepaw thus indicating functional modifications of neuronal activity. In conclusion, aperiodic spectral exponents exhibited a unique spatiotemporal profile in the mouse cortex after stroke and might complement future studies providing a dynamic link from pathophysiology to behavior.

## Introduction

Stroke is a leading cause of disability with dramatic consequences for the individual and society. Upper extremity dysfunction, for instance, is prevalent in a great portion of patients and poses major obstacles for re-integration in daily life and professional activities^1,2^. Therefore, rehabilitation after stroke is one of the major assignments in clinical neurology. In order to identify new therapeutic targets, the pathophysiology of post stroke recovery needs to be further elucidated. One aspect, that is thought to play a critical role in rehabilitation is functional reorganization of cerebral networks^3^. Electroencephalography (EEG) might be one instrument to assess these reorganization processes. In clinical practice, EEG is an easy and widely available tool to monitor brain activity and quantitative EEG measures have been linked to functional outcome after stroke^4^. Most of these EEG studies focused on changes in the periodic, oscillatory components of power spectral density (PSD) such as increases in low frequency power (delta or theta), decreases within the range of higher frequency bands (alpha, beta or gamma) or a combination of both (e.g. delta-alpha ratio)^4–8^. However, it was not until recently that the aperiodic, non-rhythmic component of neurophysiological signals gained more attention in animal and human data^9,10^. The aperiodic component defines the shape of PSD that typically shows exponentially decreasing power with increasing frequency and, accordingly, is largely dominated by a 1/f^n^ - function^9,11–14^. The exponent *n* relates to the slope of the spectrum in log-log space. The spectrum slope as well as the aperiodic exponent have been shown to be dynamic parameters in the context of cognitive states, aging and disease^9,13,15,16^. Physiologically, aperiodic exponents might reflect synaptic excitation/inhibition (E/I) balance such that increases in inhibition would lead to greater exponents^15^. Given that molecular modifications on the level of synapses have been described in the peri-infarct cortex of mice, we hypothesized that ischemic stroke would influence cortical E/I balance in its spatial vicinity which, in turn, would result in detectable changes of the aperiodic, spectral exponent^17–19^.

To that end, we recorded bilateral multichannel electrocorticography (ECoG) from widespread areas of mouse cortex in the subacute and early chronic phase after ischemic stroke and closely monitored behavioral deficits and their recovery over time. Using the permanent medial cerebral artery occlusion (pMCAO) circumscribed infarcts were induced in the sensory cortex in accordance with existing literature^20,21^. Following the procedure, affected animals showed significant impairments in a Single Pellet Reaching (SPR) task, which assesses fine motor skills of the forepaw. ECoG recordings during free exploration and immobility revealed broadband modifications of power resulting in an increased steepness of their shape particularly in adjacent cortical areas. Consequently, aperiodic exponents increased during both locomotion as well as in phases of immobility in a locally specific manner. Their normalization within the motor cortex was accompanied by a recovery of fine motor function of the contralateral forelimb hence corroborating the functional significance of this parameter.

## Results

### pMCAO induces ischemic strokes in the somatosensory cortex

All mice were implanted with a 16-channel ECoG array and individual forepaw preference was determined in a skilled reaching task. Then, ischemic stroke was induced contralaterally through permanent coagulation of the distal part of the medial cerebral artery (“stroke”: n = 15 mice). In a second cohort, identical surgical procedures were applied but the artery was left untouched (“sham”: n = 11 mice). The side of intervention was comparably distributed in the two groups (*right hemisphere, stroke: 5/15 mice, 33.33%, versus sham: 3/11 mice, 27.27%*). At the time of intervention animals of both cohorts did not differ by age (12 weeks; *82.47 ± 0.24 versus 82.09 ± 0.46 days, t(24) = 0.79, p=0.439, unpaired two-tailed t-test*) or weight (*stroke: 25.15 ± 0.19 versus sham: 25.22 ± 0.39 g, t(24) = 0.18, p = 0.859, unpaired two-tailed t-test*). Ischemic lesions almost exclusively affected the somatosensory cortex and did not expand to subcortical regions (Fig. 1a; Supplementary Tab. 1). Accordingly, numbers of cell bodies in the primary motor cortex (M1) did not differentiate between ipsi- and contralateral hemispheres of stroke and sham animals (*stroke ipsilateral: 2128 ± 49 versus stroke contralateral: 2081 ± 39 versus sham ipsilateral: 2121 ± 35 versus sham contralateral: 2144 ± 29 Nissl cells, F(3, 46) = 0.44, p = 0.726, one-way ANOVA*; Fig. 1b). Involvement of the motor cortex could not safely be excluded in one animal which was subsequently excluded from further analysis. Infarct volume in the stroke group amounted to 5.09 ± 0.97 mm^3^ on day 28, whereas in the sham group no cortical damage was observed.

**Fig. 1.**
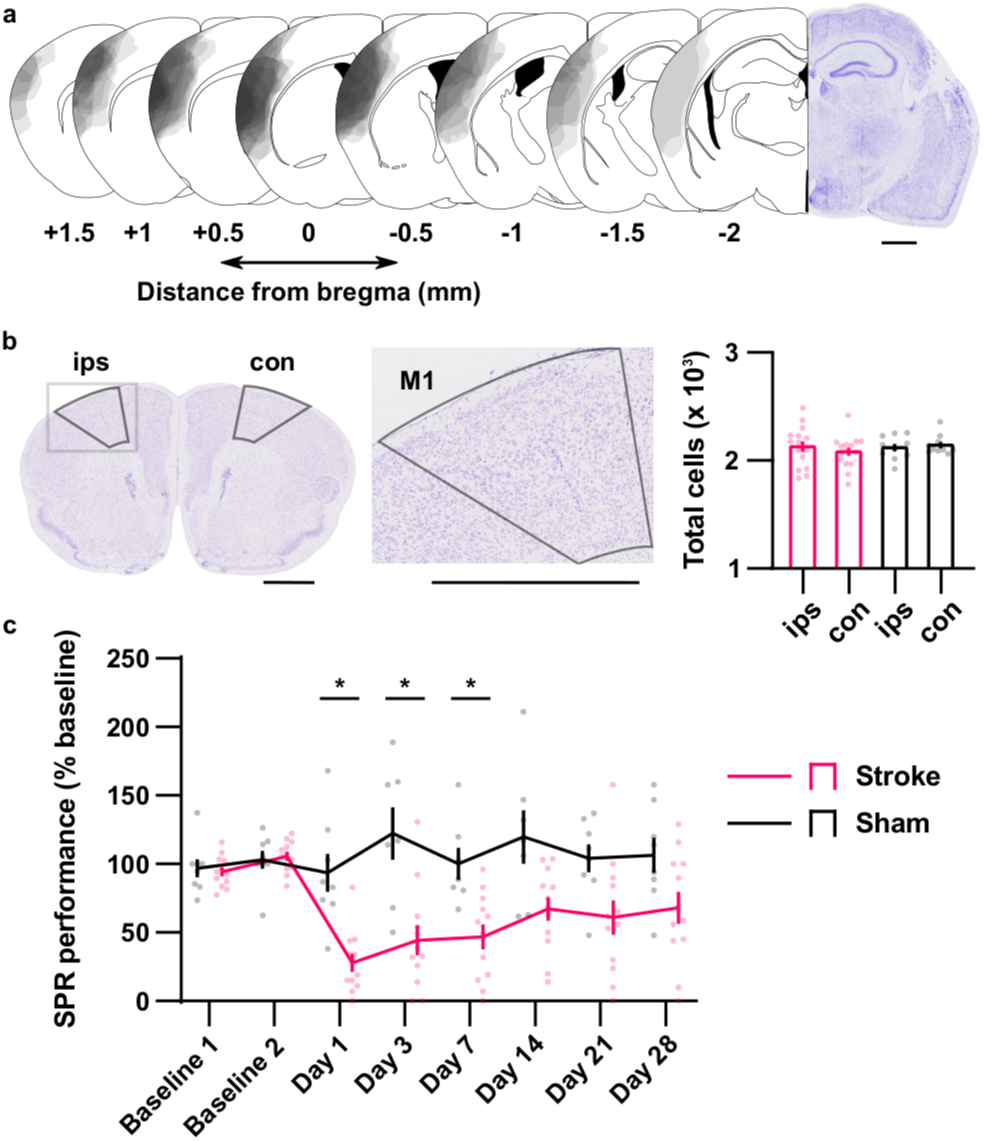
pMCAO induces fine motor deficits. **a**. Right, stroke was confirmed visually on coronal brain sections after cresyl violet staining. Left, corresponding schematics. Shaded, overlying areas indicate individual lesions. Coordinates are given in relation to bregma. **b**. Identification of M1. Cell count was performed in the area outlined. Stroke: n = 15, sham: n = 10. Ips, ipsilesional, con, contralesional. **c**. SPR performance in relation to baseline (defined as the average of the last two training sessions, baseline 1 and 2). Stroke: n = 12, sham: n = 8. Black scale bars in a and b indicate 1 mm. * p < 0.05. Data is shown ± SEM.

### Infarcts in the somatosensory cortex cause deficits in fine motor behavior

Given the wide extension of ischemic lesions within the sensory cortex, fine motor behavior such as skilled reaching might be compromised. To longitudinally assess the individual level of performance a SPR task was used, in which mice were placed inside a transparent box and were trained to reach for food pellets outside the box. 76.92% of all subjects successfully learned the task (n = 20 mice). Paw preference was equally distributed (*left forepaw preference, stroke: 4/12 mice, 25%, versus sham: 2/8 mice, 25%*). After pMCAO the percentage of successful reaches in comparison to baseline remained high in the sham group, while in the stroke group it dropped markedly to 28.00 ± 6.50% on day 1 and significantly differed from sham controls until day 7 (*intervention effect F(1, 18) = 16.93, p = 0.001, time effect F(4.328, 76.05) = 5.05, p = 0.001, intervention*time interaction F(7, 123) = 5.29, p < 0.001, repeated-measures ANOVA*; Fig. 1c). Fine motor function only partly recovered in the stroke group. After 2 weeks a performance of 67.17 ± 8.72% was reached and maintained throughout. On the individual level however, some subjects fully recovered, while others showed a beginning improvement of function before performance worsened again. Despite impairments in skilled reaching, affected animals were able to nourish sufficiently and compared to sham did not show any significant reduction of body weight throughout the course of the experiment (*intervention effect F(1, 24) = 0.59, p = 0.449, time effect F(5.889, 141.6) = 20.17, p < 0.001, intervention*time interaction F(28, 672) = 0.97, p = 0.513, repeated-measures ANOVA*; Supplementary Fig. 1b).

In contrast, mice of both groups did not present any obvious constraints of spontaneous movement after intervention surgery. Exposed to a circular Open Field arena, animals moved with a similar maximum speed (*intervention effect F(1, 24) < 0.01, p = 0.989, time effect F(6, 144) = 9.18, p < 0.001, intervention*time effect F(6, 144) = 1.04, p = 0.402, repeated-measures ANOVA*; Supplementary Fig. 2a), travelled a comparable total distance throughout the 12 minutes duration of experiment (*intervention effect F(1, 24) = 0.19, p = 0.667, time effect F(6, 144) = 7.62, p < 0.001, intervention*time interaction F(6, 144) = 0.37, p = 0.900, repeated-measures ANOVA*; Supplementary Fig. 2a) and performed ipsi- or contralateral turns equally with respect to their body axis (ipsilateral turns: *intervention effect F(1, 24) < 0.01, p = 0.964, time effect F(4.295, 130.1) = 2.59, p = 0.037, intervention*time interaction F(6, 144) = 0.87, p = 0.522*; contralateral turns: *intervention effect F(1, 24) < 0.01, p = 0.953, time effect F(3.995, 95.88) = 3.49, p = 0.011, intervention*time interaction F(6, 144) = 0.23, p = 0.966, repeated-measures ANOVA*; Supplementary Fig. 2a). All mice showed a gradual habituation to the environment as there was an increasing trend of spending time immobile over the course of 28 days (Supplementary Fig. 2a), but no differences in time spent mobile or immobile could be observed between the two groups at any time point (mobile: *intervention effect F(1, 24) = 0.02, p = 0.877, time effect F(6, 144) = 17.75, p < 0.001, intervention*time interaction F(6, 144) = 0.86, p = 0.525*; immobile: *intervention effect F(1, 24) = 0.01, p = 0.920, time effect F(6, 144) = 19.00, p < 0.001, intervention*time interaction F(6, 144) = 0.76, p = 0.60, repeated-measures ANOVA*; Supplementary Fig. 2a). Further, stroke and sham animals stayed comparably long in peripheral and central zones of the arena (periphery: *intervention effect F(1, 24) = 0.28, p = 0.599, time effect F(6, 144) = 6.89, p < 0.001, intervention*time interaction F(6, 144) = 0.85, p = 0.532*; center: *intervention effect F(1, 24) = 0.51, p = 0.483, time effectF(6, 144) = 1.34, p = 0.242, intervention*time interaction F(6, 144) = 1.14, p = 0.340, repeated-measures ANOVA*; Supplementary Fig. 2a).

Laterality of movement patterns was evaluated more specifically in the Corner and the Cylinder Test. Here, mice showed no differences in ipsi-versus contralateral turns in the Corner Test (*intervention effect F(1, 24) = 0.1.68, p = 0.207, time effect F(4, 96) = 1.16, p = 0.335, intervention*time interaction F(4, 96) = 1.57, p = 0.188, repeated-measures ANOVA*; Supplementary Fig. 2b) or in forepaw usage while exploring a transparent cylinder (*intervention effect F(1, 14) = 0.06, p = 0.811, time effect F(5, 70)= 3.09, p = 0.01, intervention*time interaction F(5, 70) = 0.97, p = 0.442, repeated-measures ANOVA*; Supplementary Fig. 2b). In summary, despite significant deficits of contralateral fine motor function no differences in locomotion or exploratory drive, and no signs of laterality of movement patterns could be detected after sensory cortical stroke.

### Aperiodic spectral exponents change in a spatiotemporal manner after pMCAO

Next, stroke-related changes in ECoG potentials were assessed. Data was recorded from freely behaving mice in a circular arena. One animal in the sham group had to be excluded from ECoG analysis due to poor signal quality (sham: n = 10 mice). Stroke and sham recordings did not differ in data length included for the spectrum analysis throughout the course of the experiment (*average data length for locomotion: 39.65 ± 1.71 s versus rest: 117.20 ± 5.57 s; locomotion: intervention effect F(1, 23) = 0.19, p = 0.671, time effect F(3.401, 78.23) = 12.86, p < 0.001, intervention*time interaction F(6, 138) = 0.17, p = 0.985, repeated-measures ANOVA; rest: intervention effect F(1, 23) = 0.03, p = 0.865, time effect F(4.334, 98.23) = 5.02, p = 0.001, intervention*time interaction F(6, 136) = 1.63, p = 0.143, mixed-effects model*). During movement and exploration ECoG recordings were dominated by rhythmic oscillations (7-8 Hz) within the theta range, during rest slower rhythms (4-5 Hz) emerged (Fig. 2; Supplementary Fig. 1c, 3a and 4). Since changes in spectral power can reflect distinct physiological processes such as true alterations of oscillatory power, shifts in the peak frequency of an oscillation, different levels of broadband power and modified spectral shapes as revealed by spectrum exponent, we evaluated periodic (oscillatory power, peak frequency) and aperiodic (exponent, offset) components of the spectra^14^. Further, due to the dependence of spectral exponents on cognitive and behavioral states, all analyses were performed separately for locomotion and rest (*baseline exponents during locomotion: 0.99 ± 0.02 versus rest: 1.42 ± 0.02, t(24) = 17.15, p < 0.001, n = 25 mice, paired two-tailed t-test*; Supplementary Fig. 1d).

**Fig. 2.**
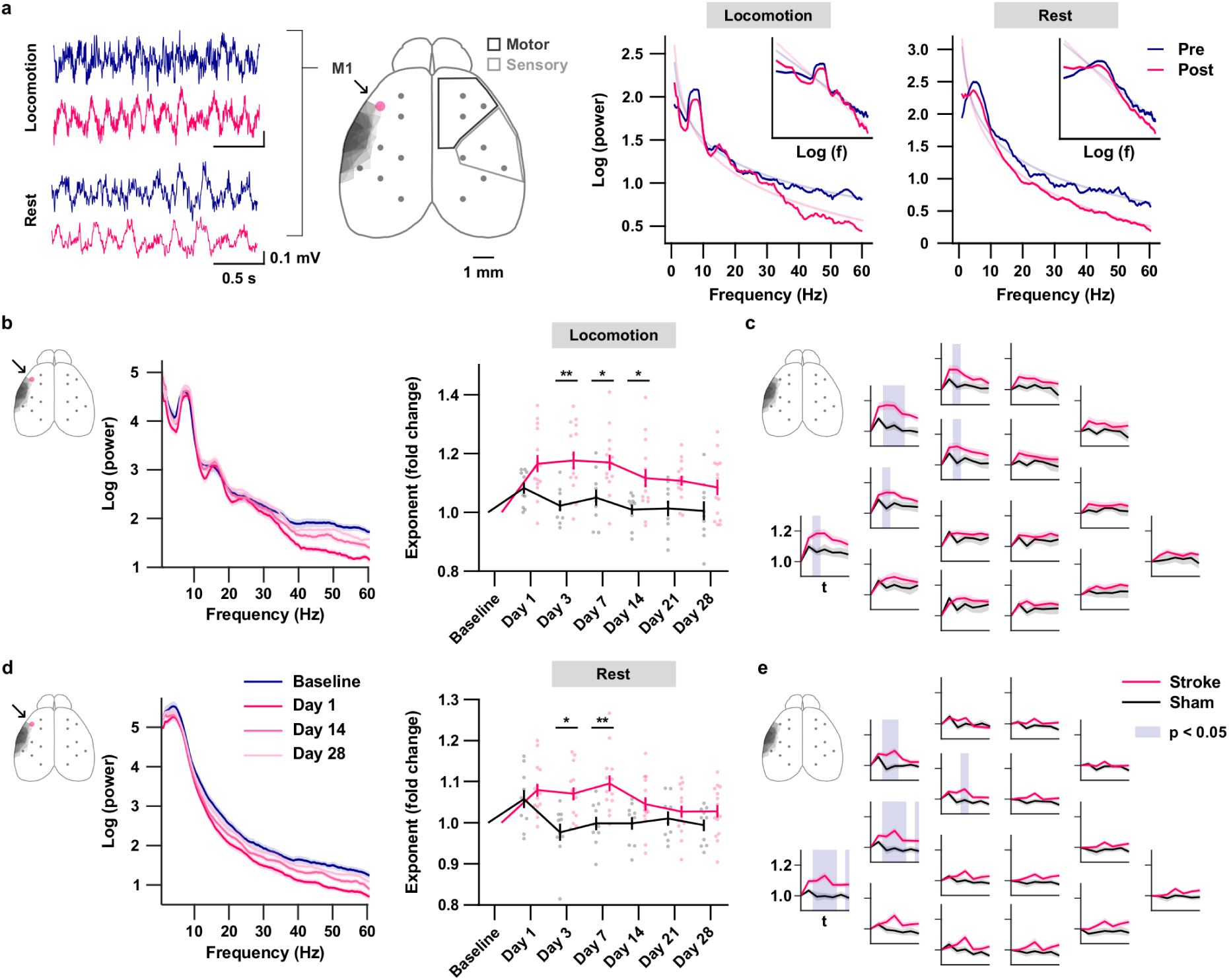
Aperiodic exponents increase in the vicinity of cortical stroke. **a**. Middle, schematic illustrating individual strokes (shaded areas) in relation to electrode position and motor and sensory cortical areas. Red dot indicates channel 3 covering M1. Left, ECoG data of a single subject recorded from M1 (black arrow, red dot) before (blue) and after (red) stroke induction are shown. Right, corresponding PSD from M1 of the same subject in semi-log and log-log space (inset). Transparent lines correspond to regression lines in log-log space. F, frequency. **b**. Left, semi-log plots of group-averaged PSD of M1 (black arrow, red dot) over time before and after stroke induction during locomotion. Right, corresponding time course of aperiodic exponents (red) compared to sham controls (black). The fold change of aperiodic exponents compared to their baseline values is presented. **c**. Group-averaged overview of time courses of aperiodic exponents for all 16 ECoG channels recorded during locomotion. The fold change of aperiodic exponents compared to their baseline values is presented. Shaded areas indicate significant differences. **d**. and **e**. Same as b. and c. but during rest. Stroke: n = 15, sham: n = 10. ** p < 0.01, * p < 0.05. Data is shown ± SEM.

After stroke induction, a broadband loss of power could be observed in PSD in the vicinity of the ischemic lesion (Fig. 2a). Since the decrease of power seemed to augment with increasing frequency, this loss caused a tilt of the entire corresponding spectra. Subsequently in M1, spectrum slope appeared steeper after stroke (Fig. 2a). Accordingly, aperiodic exponents increased for both behavioral conditions (Fig. 2b and d) and remained significantly different from sham controls until day 14 during locomotion (*intervention effect F(1, 23) = 12.52, p = 0.002, time effect F(3.617, 83.19) = 9.95, p < 0.001, intervention*time interaction F(6, 138) = 3.08, p = 0.007, repeated-measures ANOVA*; Fig. 2b) and until day 7 during rest (*intervention effect F(1, 23) = 5.93, p = 0.023, time effect F(3.951, 89.56) = 9.94, p < 0.001, intervention*time interaction F(6, 136) = 6.35, p < 0.001, mixed-effects model*; Fig. 2d). Considering all recorded channels during locomotion, significant alterations of aperiodic exponents could be observed particularly long in channels covering the ipsilesional motor cortex, while during rest channels close to ipsilesional sensory areas were more severely affected (Fig. 2c and e). Broadband power (1-60 Hz) significantly differed from sham controls only during rest in the most anterior channels covering parts of the motor cortex (Supplementary Fig. 3b) and aperiodic offset did not reveal significant differences (Supplementary Fig. 3c).

Focusing on rhythmic oscillatory activity, within the motor cortex three spectrum peaks could be identified at 7-8 Hz and 15-16 Hz during locomotion, and 4-5 Hz during rest. Following pMCAO, their relative power did not differ between the two groups regardless of the behavioral condition considered (locomotion, power peak 1: *intervention effect F(1, 23) = 1.65, p = 0.212, time effect F(4.410, 101.4) = 2.83, p = 0.025, intervention*time interaction F(6, 138) = 0.96, p = 0.453, repeated-measures ANOVA*; locomotion, power peak 2: *intervention effect F(1, 23) = 0.60, p = 0.447, time effect F(4.963, 114.2) = 1.39, p = 0.232, intervention*time interaction F(6, 138) = 1.00, p = 0.430, repeated-measures ANOVA*; rest, power peak 3: *intervention effect F(1, 23) = 1.19, p = 0.287, time effect F(4.737, 107.4) = 1.92, p = 0.101, intervention*time interaction F(6, 136) = 1.03, p = 0.408, mixed-effects model*; Supplementary Fig. 4a and b). Peak frequency remained constant (locomotion, frequency peak 1: *intervention effect F(1, 23) = 3.85, p = 0.062, time effect F(4.362, 100.3) = 3.94, p = 0.004, intervention*time interaction F(6, 138) = 2.86, p = 0.012, repeated-measures ANOVA*; locomotion, frequency peak 2: *intervention effect F(1, 23) = 0.10, p = 0.753, time effect F(3.879, 73.05) = 2.20, p = 0.079, intervention*time interaction F(6, 113) = 0.70, p = 0.649, mixed-effects model*; rest, frequency peak 3: *intervention effect F(1, 23) = 0.62, p = 0.439, time effect F(3.769, 65.33) = 1.06, p = 0.380, intervention*time interaction F(6, 104) = 2.15, p = 0.054, mixed-effects model*; Supplementary Fig. 4a and b). In summary, sensory cortical stroke induced widespread alterations in the aperiodic, but not the periodic components of recorded ECoG. These alterations could be observed in the motor cortex even though being histologically intact.

### Spectrum shape correlates with behavioral deficits

To evaluate if aperiodic exponents hold functional significance, temporal correlations between electrophysiological and behavioral metrics each measured on the same day after stroke induction were investigated. During locomotion, aperiodic exponents and SPR performance rate were significantly correlated in ipsilateral motor (*M1, channel 3: R*_*rm*_ *= -0.36, p = 0.005*) and adjacent areas (Fig. 3a). Contralateral motor cortex showed significant correlations to a lesser extent (Fig. 3a). During rest, correlations were more strictly limited to the ipsilateral motor cortex (*M1, channel 3: R*_*rm*_ *= -0.27, p = 0.034*; Fig. 3b). Next, to investigate whether correlations were present due to power changes within circumscribed frequency bands, power spectra were segmented into frequency bins of 1 Hz and correlations of each bin with SPR performance rate were determined. Within motor cortex significant correlations were detected for most frequencies > 17 Hz during movement and almost the entire spectrum during rest (Fig. 3c). No particular frequency bin exceeded regarding the calculated R_rm_-coefficients. Thus, behavioral performance did not seem to solely depend on power changes within specific frequency bands but rather on broadband modifications of spectra that might be evoked by alterations of aperiodic exponents. Stroke volume did not correlate with SPR performance, with spectral exponents or with broadband power in the motor cortex (Supplementary Tab. 2).

**Fig. 3.**
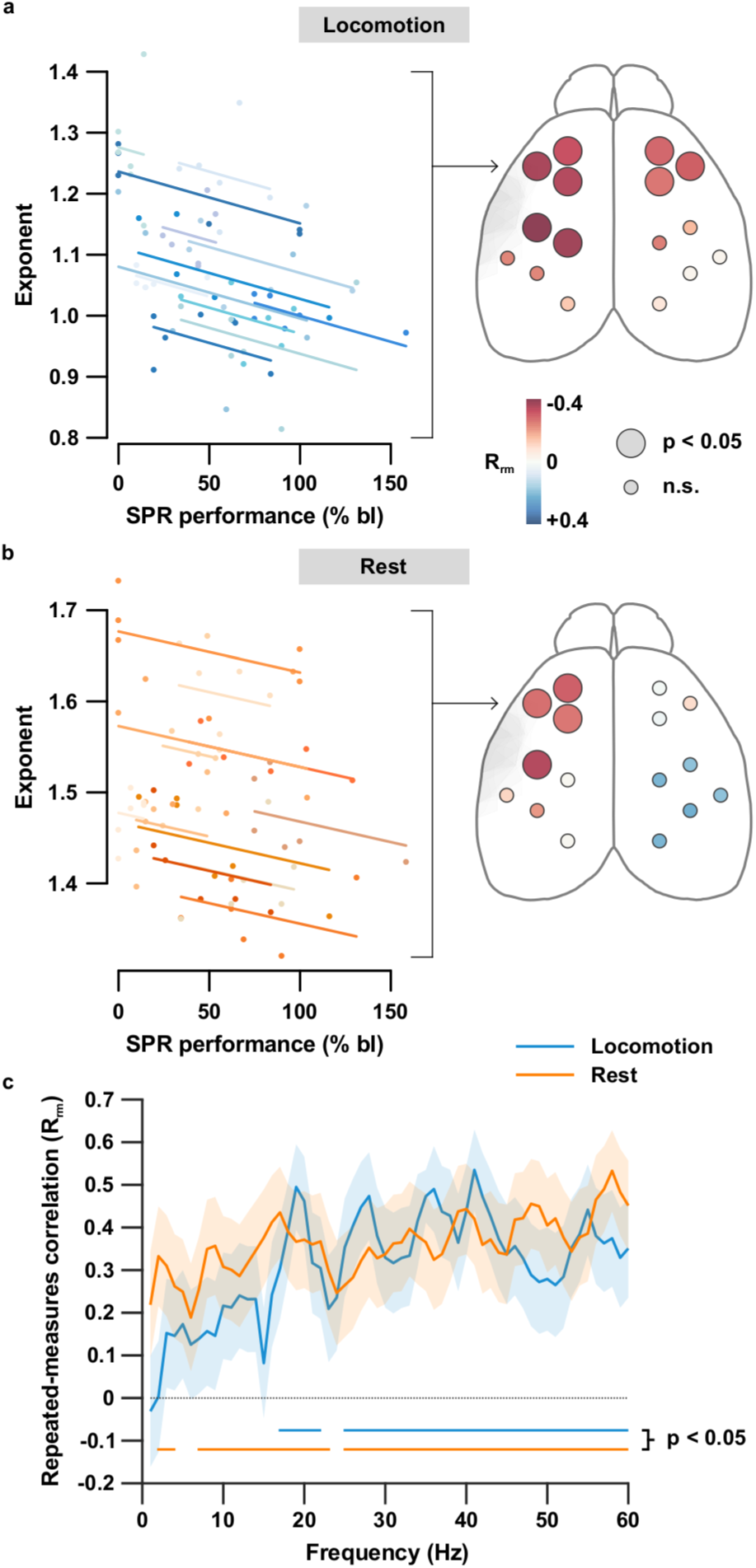
Exponents from motor areas correlate with behavioral performance. **a**. During locomotion a negative correlation between aperiodic exponents and SPR performance after ischemia is revealed through repeated-measures correlation. Left, exemplary plot depicting correlation for M1 (arrow). Dots and lines of the same color represent values of individual animals. Right, schematic illustrating subsequent correlation values (R_rm_) for all 16 channels (color-coded). Large dots indicate significant correlations. **b**. Same as a., shown for the resting condition. **c**. Repeated-measures correlation values between SPR performance and power of single frequency bins of 1 Hz each during locomotion (blue) and rest (orange). Blue and orange lines on the bottom indicate significant correlations. n = 12. Data is shown ± SEM.

## Discussion

This study characterized transient increases of spectral, aperiodic exponents in multichannel ECoG data of mice after sensory cortical stroke. While individual lesion size did not shape the evaluated electrophysiological and behavioral parameters of an animal, exponent normalization in the motor cortex matched the time course of fine motor recovery of the contralateral forepaw.

In order to realize complex movements such as skilled reaching the primary motor cortex might depend on unimpeded sensorimotor integration of information that comprises proprioceptive feedback originating from primary or secondary sensory areas^22,23^. In human patients, ischemic lesions within sensory cortical areas impair motor recovery of the hand^24^. Here, not the structural extent of infarcts but functional alterations in the vicinity, that could be tracked through ECoG, correlated with functional deficits. PSD analysis of ECoG recordings revealed a much steeper spectrum shape shortly after intervention that manifested in increased aperiodic exponents. During recovery, exponents returned to baseline levels closely following the time course of fine motor recovery which resulted in a significant correlation. In contrast, power within well described frequency bands such as theta (7-8 Hz) during locomotion or respiration-related oscillations (4-5 Hz) during rest remained stable^25–27^. The absolute value of correlation between aperiodic exponents and behavior, however, was negatively influenced by the observation that in the reaching task single mice showed a distinct recovery after a few days which was then not maintained or even reverted throughout the time course. A possible reason might be, first, a refinement of the SPR protocol, in which, in order to reduce the animals’ burden, food restriction was only applied in nights prior to an experiment and not constantly as suggested elsewhere^28^. Therefore, some subjects might have lost motivation to perform as soon as they learned that regular food would be returned sometime after the test. Second, sensory deafferentation with preserved motor function could have led to a reduced utilization of the affected forepaw after previous unsuccessful attempts, a phenomenon termed ‘learned nonuse’ which imposes negative consequences for cortical reorganization in the context of stroke rehabilitation and which also could have interfered with the results^29,30^. Following sensory cortical ischemia, locomotion was not impaired in affected animals in this study since being a relatively primitive motor skill that, even in the absence of cortical control, might be maintained through specialized central pattern generators in the spinal cord of mice^31^. Further, no signs of laterality in movement patterns could be observed in the Open Field, Corner or Cylinder Tests in accordance with previous studies^32^. In summary, these results indicate that potentially reversible, functional modifications of cortical activity, to some extent, determine behavioral outcome after pMCAO.

Spectral exponents or slopes have been interpreted to reflect E/I balance. Gao and colleagues (2017) delivered a computational model that assumes a high conductance state of cortical circuits such as in the absence of stimulus-specific responses and a linear summation of excitatory and inhibitory currents with fast glutamate and slower GABA signaling, respectively. In such a model, spectral slopes would appear steeper when E/I balance shifts towards inhibition and vice versa^15^. *In vivo* evidence exist in the context of various anatomical and neurophysiological mechanisms where, for instance, spectrum slope reflects the spatial gradients of excitatory versus inhibitory synapse densities along dendrites in rat hippocampus, the rhythmicity of theta-modulated excitatory and inhibitory time windows, and propofol-induced and GABA-mediated inhibition in anesthesia^15^. In addition, the functional relevance of spectral slopes or exponents has been demonstrated in relation to age, cognitive tasks such as working memory, and disease^13,14,16^. Considering the presented data, ECoG consists of a summation of postsynaptic currents and consequently excitatory and inhibitory synaptic currents must also be reflected^15,33^. A model in which E/I balance drives spectrum shape allows an interpretation where an increased spectral steepness would indicate heightened levels of inhibition. Clarkson and colleagues (2010) found increased tonic inhibition in perilesional cortex due to impaired GABA transport. Corresponding behavioral deficits could be attenuated by blocking GABA-mediated signaling^17^. Similarly, in the present study increased spectral exponents were found predominantly in adjacent ipsilateral areas and in the motor cortex. A normalization of exponents was accompanied by a recovery of fine motor function. Thus, a temporary shift of E/I balance towards inhibition in the respective areas might have impeded behavioral recovery. The underlying pathophysiology, however, needs to be further clarified in future studies disentangling the molecular pathways of structural and functional reorganization of neural activity.

In the past, quantitative EEG (qEEG) measures in stroke research have mainly focused on changes in oscillatory power. Power asymmetry between ipsi-and contralateral hemispheres has been proposed in form of a brain symmetry index (BSI) to indicate unilateral ischemia and estimate clinical prognosis in human patients^34,35^. Delta power was used in a similar manner as acute delta change index (aDCI) or in relation to alpha power as delta/alpha-ratio (DAR)^36,37^. However, in an event where a spectrum is tilted and in which broadband changes with varying impact on circumscribed frequency bands occur, these measures might yield ambiguous results. A recently published study in mice showed a decrease of power in the gamma frequency band (30-50 Hz) in local field potentials of the perilesional cortex after photothrombotic stroke^38^. This finding was apparent when compared to the contralateral hemisphere but not in comparison to sham animals^38^. Although aperiodic components were not specifically evaluated in that study, they might have contributed to the results since power reduction did not exclusively affect gamma but was accompanied by changes in several other frequency bands^38^. Electrophysiological observations in mice after stroke are not unlikely to occur in humans^39,40^. Spectral exponents in human stroke patients are under investigation^10^. A joint analysis of periodic and aperiodic parameters might enable future studies to further reveal their interference and prognostic value in stroke rehabilitation.

There are limitations to our study. In contrast to previous work, increased levels of slow frequency power were absent. Slow synchronous neuronal activity has been observed after cortical stroke and has been linked to axonal sprouting originating from the contralateral homotypic cortex^41^. Further, slow-wave activity has been described close to the ischemic lesion possibly reflecting hyperpolarization and inhibition of cortical neurons or, alternatively, a default-mode of deafferentiated, cortical neurons^37,42,43^. The analysis of slow-wave activity was not the focus of this study and might have been methodologically compromised by the 2s-length of data episodes resulting in a Rayleigh frequency of 0.5 Hz. This was specifically chosen because of the short-lived, spontaneous episodes of behavior exhibited by freely behaving mice. Further, due to the experimental design recordings were performed during daytime for fixed durations and not continuously as in other studies^9,44^. Interestingly, some animals showed a transient increase of exponents after sham surgery which, therefore, did not differ significantly from stroke on day 1. It might be possible that the opening of the skull during the intervention had set a stimulus with a transient impact on neuronal function. If sham surgery evokes any pathological response needs to be further studied. In both groups an increase of broadband power could be observed over time. Due to the chronic character of the experiment, even though electrode implants did not permeate the dura, glial scarring and meningeal lymphangiogenesis might have been induced forming a resistive layer, that could possibly affect the impedance and, thus, the signal-to-noise ratio of an electrode^45,46^. Yet, it should be highlighted that this aspect did not manifest in changes of spectral exponents since they remained stable in the sham group throughout the course of the experiment.

In conclusion, this is the first study that systematically employs bihemispherical, multichannel ECoG in mice after ischemic stroke. Aperiodic exponents which are considered to reflect cortical E/I balance changed in a locally specific manner after cortical ischemia and their normalization within the motor cortex was accompanied by a recovery of fine motor function of the contralateral forelimb. The findings of this study contribute to the understanding of post stroke pathophysiology and might facilitate the identification of spatial and temporal windows for future innovative treatment approaches. Since mice are widely used to study the mechanistic foundation of stroke recovery, the analysis of aperiodic components in electrophysiological signals might help to close the gap between behavioral readouts and the activity state of neuronal populations.

## Methods

### Ethics statement

C57/Bl6 male mice aged nine postnatal weeks at the start of the experiment were used in this study. Experimental protocols were approved by the Behörde für Justiz und Verbraucherschutz der Freien und Hansestadt Hamburg (approval number N53/2020). All procedures were performed under the guidelines of the animal facility of the University Medical Center Hamburg-Eppendorf and in compliance with the Guide for the Care and Use of Laboratory Animals. Animals were housed in groups under a 12 h dark-light cycle. After implantation of electrodes, mice were housed individually for 49 days.

### Electrode implantation and pMCAO surgery

ECoG electrodes were chronically implanted. Prior to implantation surgery, mice received a subcutaneous injection of Buprenorphine (0.05 mg/kg body weight). Then, animals were anaesthetized with isoflurane (induction 3-5%, maintenance 1-2% in O2 at an airflow of 0.5 l/min) and fixed in a stereotaxic frame. Body temperature was kept stable with a heating pad. 16 craniotomies (0.5 - 1 mm diameter) were performed to implant 16 perfluoroalkoxy alkane coated platinum/iridium wire electrodes of 127 μm diameter (101R-5T, Science Products) just above the cortex of both hemispheres leaving the dura mater intact according to stereotaxic coordinates (Supplementary Tab. 3)^47^. A reference consisting of the same material was placed over the cerebellum, a stainless-steel screw (1.2 mm diameter) served as ground. An additional screw placed over the cerebellum was used to further stabilize the implant. All electrodes were connected to a connector (A79038-001, Omnetics). A drop of superglue was used to cover the craniotomies and stabilize the electrodes before dental cement was used to finally fix the electrodes in place. For pMCAO surgery, anesthesia was performed in the same way as described above. Importantly, ischemic stroke was induced on the hemisphere contralateral to the previously determined, dominant forepaw of an animal. If a subject did not show any paw preference the hemisphere was randomly chosen. The following procedure was described in detail previously^48^. In short, mice were put in a lateral position on a heating pad, a skin incision of 1 cm was made between the eye and ear, the temporal muscle was cut with a coagulation forceps and flapped away. Then, the bifurcation of the medial cerebral artery (MCA) was identified and a small craniotomy was performed. Now, all three branches of the bifurcation were coagulated with finer coagulation forceps. If no bifurcation could be identified the most proximal part of the MCA was coagulated. In sham controls the surgery was performed the same way except for the coagulation of the MCA. After both electrode implantation and pMCAO or sham surgery mice received water and soft food to which tramadol was added for 72 hours. Before any behavioral training or electrophysiological recording started mice were granted a recovery period of one week after electrode implantation.

### *In vivo* electrophysiology

ECoG data was recorded from freely behaving mice with a wireless amplifier system (W2100 System, HS16 Headstage, Multichannel Systems) at a sampling rate of 1000 Hz. While being recorded mice were able to freely explore a circular arena of 60 cm diameter for 12-25 minutes. The weight of the headstage and associated battery amounted to 4.9 g. At least one day before first baseline recordings took place all mice were already exposed to the arena and the headstage for acclimatization. Behavior was monitored with an overhead camera (UI-3240-NIR-GL Rev.2, IDS Imaging; W2100-Video-System-NIR, Multichannel Systems), corresponding videos were analyzed offline and episodes of locomotion and immobility (from now on termed “rest” episodes) were visually identified. Episodes with a minimum duration of 2s were used for electrophysiological data analysis. Synchrony was further guaranteed aligning electrophysiological data and video by manually triggered events (Arduino Uno, Arduino).

### Data analysis

Data analysis was predominantly done in Matlab (R2018b, The MathWorks Inc.) using the FieldTrip toolbox^49^. First, continuous data was filtered between 0.3 Hz and 180 Hz using a 4^th^-order Butterworth filter. Next, episodes of 2 s length belonging to either one of the two previously defined behavioral conditions were extracted. Data was visually inspected and channels containing artifacts throughout the recording were excluded. When artifacts occurred in single episodes, those were rejected from all channels. If the number of remaining episodes after artifact rejection was insufficient for further analysis, the corresponding recording was excluded. PSD was computed on single episodes and then averaged per behavioral condition. To that end, data was demeaned first, then Fast-Fourier Transformation after multi-tapering with 7 tapers was applied resulting in a spectral smoothing of ± 2 Hz given the 2 s length of episodes^50^. To estimate aperiodic parameters of spectra, we fitted a linear approximation to the PSD graph in log-log space between 1 and 60 Hz using a recently released toolbox (FOOOF 1.0.0) in Python 3.8.3 Spyder IDE^14^. Spectral peaks were identified using the ‘findpeaks’-function in MATLAB. Relative power was calculated dividing the power of identified peaks by overall power of the spectrum between 1 and 60 Hz. If no peak was present in a spectrum, no frequency value was extracted and relative power was estimated in between 7.5 to 8.5 and 15.5 to 16.5 Hz during locomotion and in between 4 to 5 Hz during rest corresponding to peak frequencies with strongest power on the group level. All analyses on the group level were performed after matching data from corresponding channels of either the ‘ipsilesional’ or ‘contralesional’ hemisphere. For visualization, ipsilesional channels are plotted on the left.

### Behavior

All behavioral experiments were carried out in a chamber with constant light source and video recorded for eventual verification of test scores. Mice were placed inside the chamber 30 minutes prior to an experiment for acclimatization. In between experiments the chamber as well as the respective test apparatus were wiped with 70% ethanol. Up to two different tests per mice were performed per day (Supplementary Fig. 1a).

### Open Field

General locomotor activity was assessed in an open field scenario. Mice were allowed to freely explore a circular arena of 60 cm diameter while electrophysiological data was recorded at the same time using a wireless system. Experiments consisting of 12 minutes each were recorded with an overhead camera and were analyzed offline using a commercially available software (ANY-maze Video Tracking System 6.35, Stoelting Co.). To carefully connect the headstage to the implant, mice were shortly sedated with Isoflurane prior to being put inside the chamber for acclimatization.

### Corner Test and Cylinder Test

Laterality was tested using both a Corner and a Cylinder Test. For the corner test two plastic boards of 30 × 20 cm size were attached to each other at an angle of 30°. Mice were placed half way inside and facing the corner. Trials were counted if mice approached the corner before making a turn and dismissed if the turn was made immediately. Asymmetry in usage of the forepaw was further assessed using the cylinder test. Animals were placed inside a transparent acrylic glass cylinder of 20.5 cm height and 12 cm diameter and were recorded for 5 minutes. For exploration of the cylinder mice repeatedly reared up, touched the cylinder walls with their forepaws, subsequently dismounted the respective paws and landed back on the ground. Asymmetry was assessed by counting always the first contact of a forelimb with the cylinder wall at full rear when mice stood only on their hind legs with both forelimbs off the ground^48^. If the first contact was realized with both paws simultaneously both were counted. Analysis was performed offline until at least 20 contacts for one forelimb were counted. Turns in the corner and forepaw usage in the Cylinder Test were termed ‘ipsilateral’ and ‘contralateral’ with respect to the side of stroke or sham surgery.

### Single Pellet Reaching

Fine motor skills were evaluated using a single-pellet reaching task as described previously^28^. In short, experiments were performed in a transparent acrylic glass box (20 × 15 × 8.5 cm) with 0.5 cm wall thickness that contained narrow windows of 13 cm height and 0.5 cm diameter. Mice were trained to use their forepaws to reach through a window and retrieve a food pellet placed on a shelf outside the box. In our protocol millet seeds served as food pellets. Training started one week after electrode implantation and was continued for a maximum of 14 days. Each training session lasted up to 20 minutes or until 30 reaching attempts were performed. In the first week, the dominant forepaw was determined. Therefore, pellets were placed at an easy reach until an animal presented paw-preference by making > 65% of reaching attempts within one trial with a single paw. Then, millet seeds were presented only on the preferred side of each mouse and the level of individual performance was determined. An attempt was considered successful if a mouse was able to grab a seed and eat it. In a failed attempt, mice reached for a seed but either knocked it off the shelf or dropped it to the floor. If an animal was not able to accomplish 9 successful attempts out of 30 reaches after 7 days of training it was labeled ‘non-learner’ and excluded from the analysis. The average number of successful reaching attempts on the last two days of training was defined as the individual baseline performance of a mouse. Testing sessions after intervention were performed accordingly and performance was assessed in relation to baseline (in %). If less than 30 reaching attempts within 20 minutes were completed, a trial was classified as ‘did not perform’ and not considered for further analysis. In contrast to the original protocol, access to regular food was only restricted in the night prior to an experiment and returned afterwards. Experiments were performed at fixed hours in the morning.

### Histology

After completion of the last experiment animals were terminally anesthetized with ketamine/xylazine (120 and 16 mg/kg respectively) and intracardially perfused with 20 ml of 0.1 M phosphate-buffered saline (PBS), followed by 20 ml of 4% paraformaldehyde (PFA) in PBS at pH 7.4. Brains were removed, post fixated in 4% PFA, rinsed in PBS for 24h at 4°C, dehydrated in a graded series of ethanol, embedded into paraffin blocks and coronally sectioned (5 μm slice thickness) with a microtome. Sections were then stained with 0.1% cresyl violet and photographed (Nano Zoomer 2.0-HT, Hamamatsu Photonics K.K.). Infarct size was calculated on the scanned image by tracing the area containing the ischemic neuronal damage, which was outlined based on the absence of Nissl stained neurons in the cortex using NDP.view2 software (Hamamatsu Photonics K.K.). Lesion volume was determined by multiplying the correspondent area by the section interval thickness^51^. Neuron loss in the motor cortex was assessed by comparing ipsi-and contralesional hemispheres. The number of Nissl cell bodies was estimated using the ImageJ particle analyzer plugin^52,53^.

### Statistics

Statistics were performed in Prism (9.3.0, GraphPad) and R (4.1.1, RStudio). Single comparisons were carried out using two-tailed student’s t-test. For multiple comparisons one-way ANOVA after Bonferroni correction for multiple comparisons was used. To assess effects of intervention (stroke or sham surgery) and time on behavioral performance or electrophysiological parameters two-way repeated-measures ANOVA was evaluated with Geisser-Greenhouse correction since no sphericity of the data was assumed and after applying Bonferroni correction for multiple comparisons. Since repeated-measures ANOVA cannot handle missing values in the case of exclusion of single experiments a mixed-effects model as implemented in Prism 9.3.0 that uses a compound symmetry covariance matrix was fit using Restricted Maximum Likelihood. In the absence of missing values, this method gives the same p-values and multiple comparisons tests as repeated-measures ANOVA. In the presence of missing values (missing completely at random), the results can be interpreted like repeated measures ANOVA. For correlation of infarct size with behavioral and electrophysiological parameters Spearman correlation was carried out. When single subjects provided more than one data point, repeated-measures correlation was performed in R using the rmcorr-package (0.4.4)^54^. Data is presented as mean ± SEM and was considered statistically significant when p < 0.05.

## Acknowledgements

We thank Mattia Chini for valuable discussions on the data and feedback, and Christian Hagel for providing scans of sectioned brains. This work was funded by the German Research Foundation (DFG, FOR 2879, project A3), and by the German Research Foundation and the Natural Science Foundation of China (DFG and NSFC, SFB/TRR 169, project A3).

## Author contributions

J.B. designed the study, performed experiments, analyzed the data and wrote the manuscript. S.I.C. performed experiments and analyzed the data. F.L.H. analyzed the data. T.M. designed the study and wrote the manuscript. All authors interpreted the data, discussed and commented on the manuscript.

## Corresponding author

Correspondence and requests for materials should be addressed to Jonatan Biskamp.

## Supplementary Figures

**Supplementary Fig. 1.**
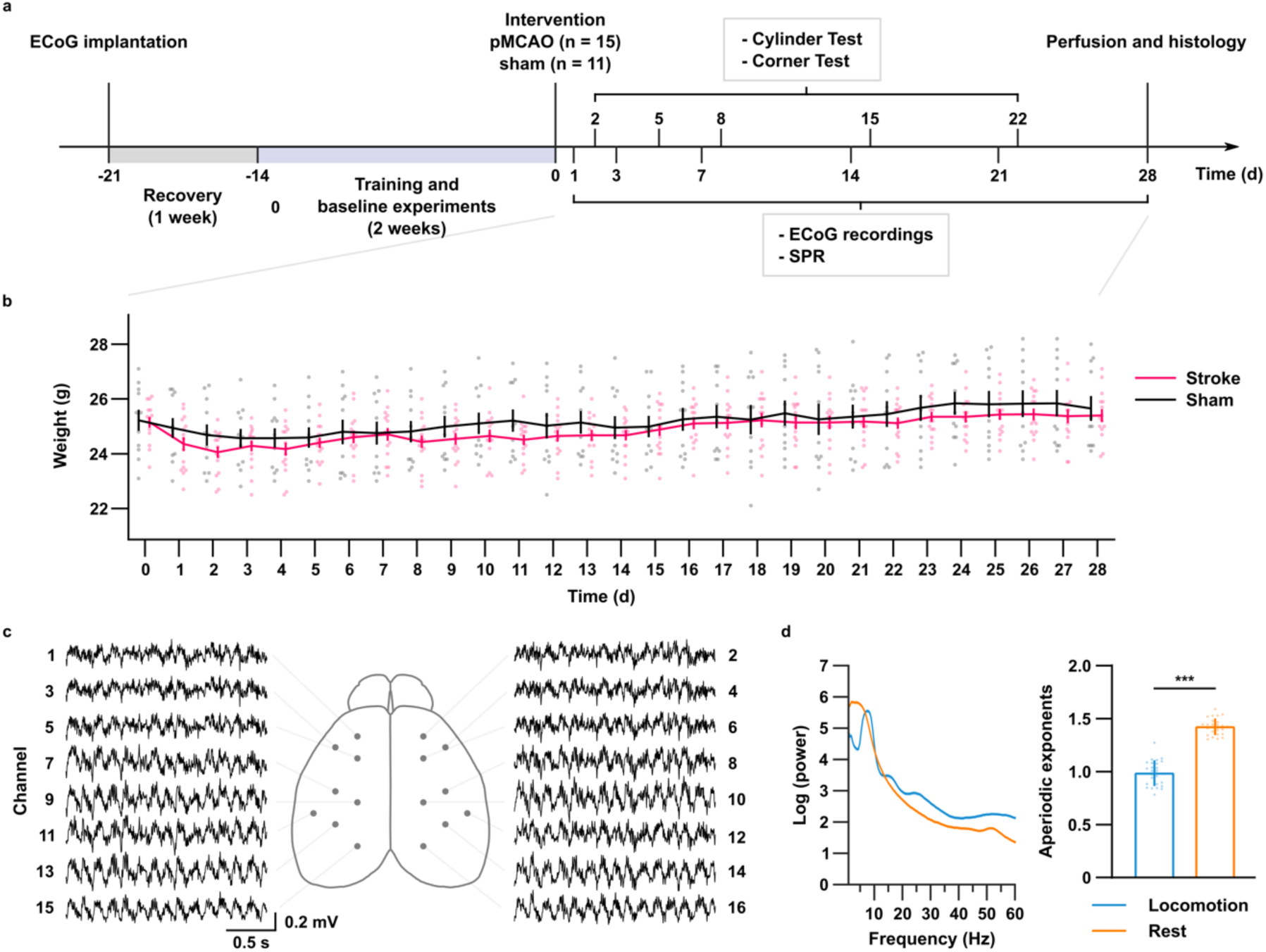
Experimental schematic and ECoG setup. **a**. Schematic illustrating time course of the experiment. **b**. Body weight before (d = 0) and after intervention for stroke (red) and sham (black) animals. D, days. Stroke: n = 15, sham: n = 11. **c**. Schematic showing electrode setup and exemplary raw ECoG data of 2 s each. By definition, odd channel numbers indicate the hemisphere of intervention. **d**. Left, semi-log plot of averaged PSD of all channels and animals during locomotion (blue) and rest (orange) before intervention. Right, corresponding aperiodic exponents. n = 25. *** p < 0.001. Data is shown ± SEM.

**Supplementary Fig. 2.**
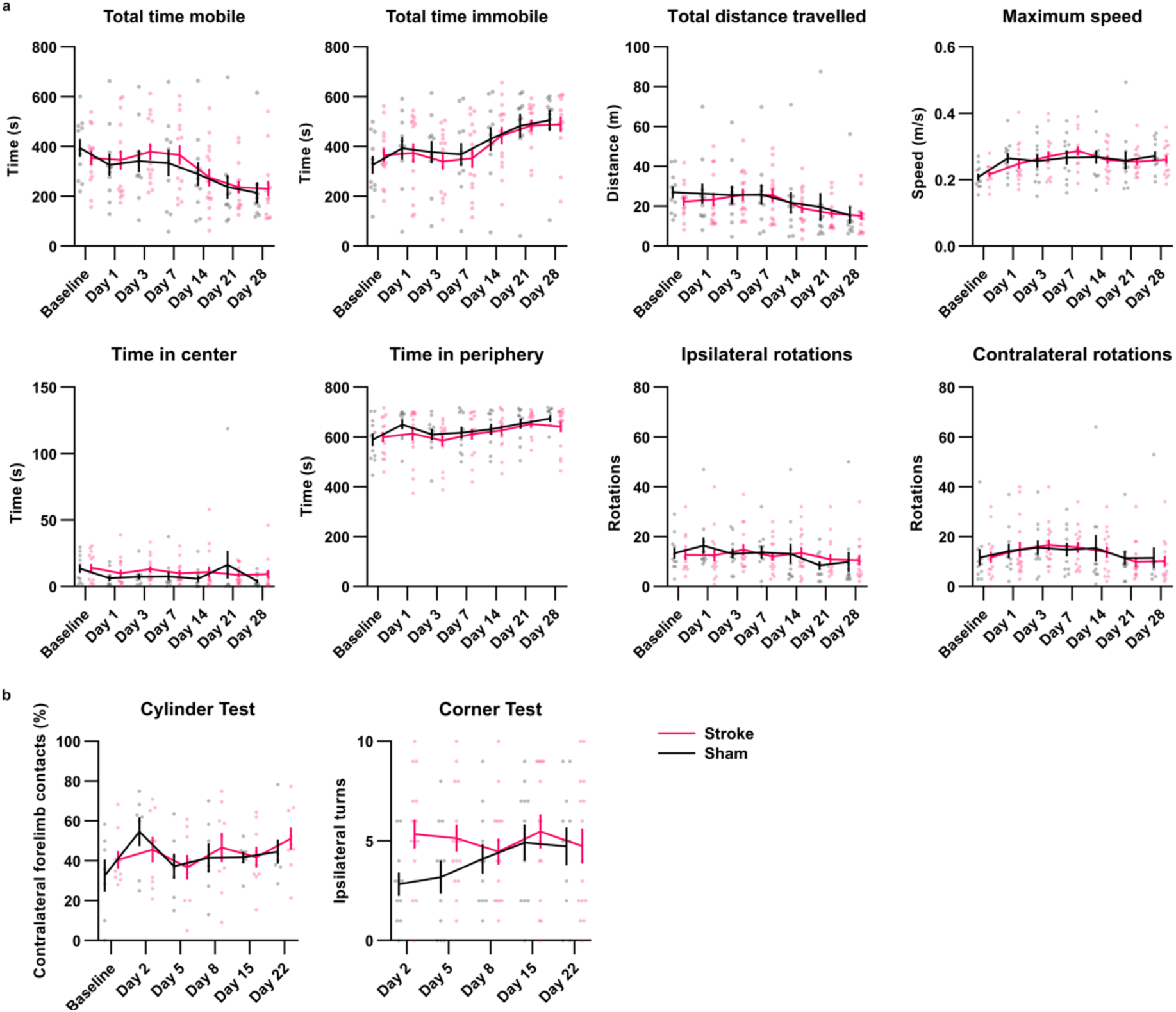
Behavioral results after sensory stroke. **a**. Plots illustrating the indicated measures while stroke (red) and sham (black) mice freely explored a circular Open Field arena for 12 minutes. **b**. Contralateral forelimb contacts and ipsilateral turns in the Cylinder and Corner Test respectively. Stroke: n = 15, sham: n= 11. Data is shown ± SEM.

**Supplementary Fig. 3.**
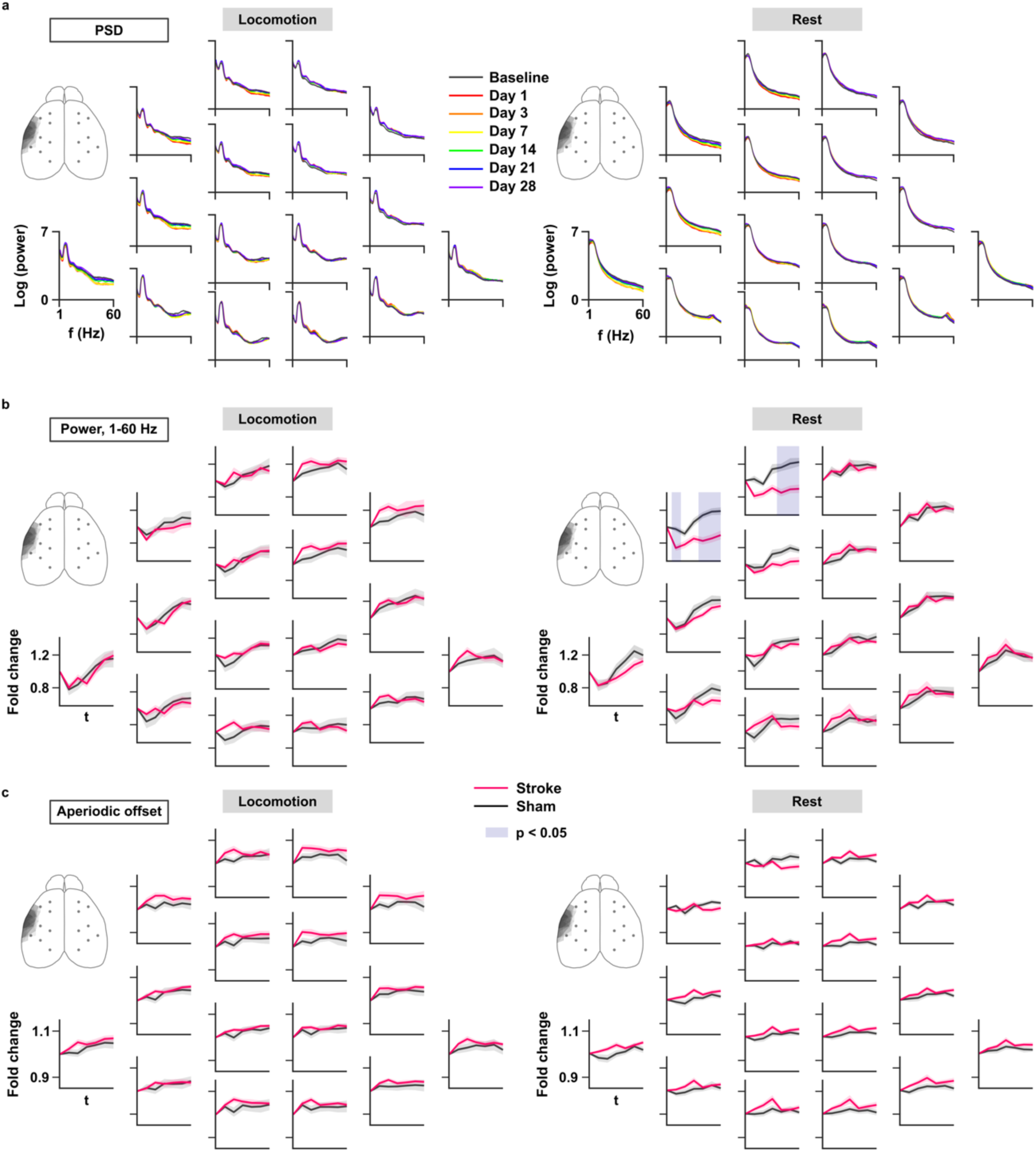
Time course of PSD changes, broadband power and spectrum offset. **a**. Overview showing semi-log plots of group-averaged PSD of all 16 channels before and after stroke during locomotion (left) and rest (right). Each color represents a different time point. F, frequency. **b**. Group-averaged overview of time courses of broadband power (1-60 Hz) for all 16 ECoG channels recorded during locomotion (left) and rest (right). Fold change in relation to baseline is shown. Shaded areas indicate significant differences between stroke (red) and sham (black) animals. T, time. **c**. Group-averaged overview of time courses of aperiodic offsets for all 16 channels during locomotion (left) and rest (right). Fold change in relation to baseline is shown for stroke (red) and sham (black) animals. Stroke: n = 15, sham: n = 10. Data is shown ± SEM.

**Supplementary Fig. 4.**
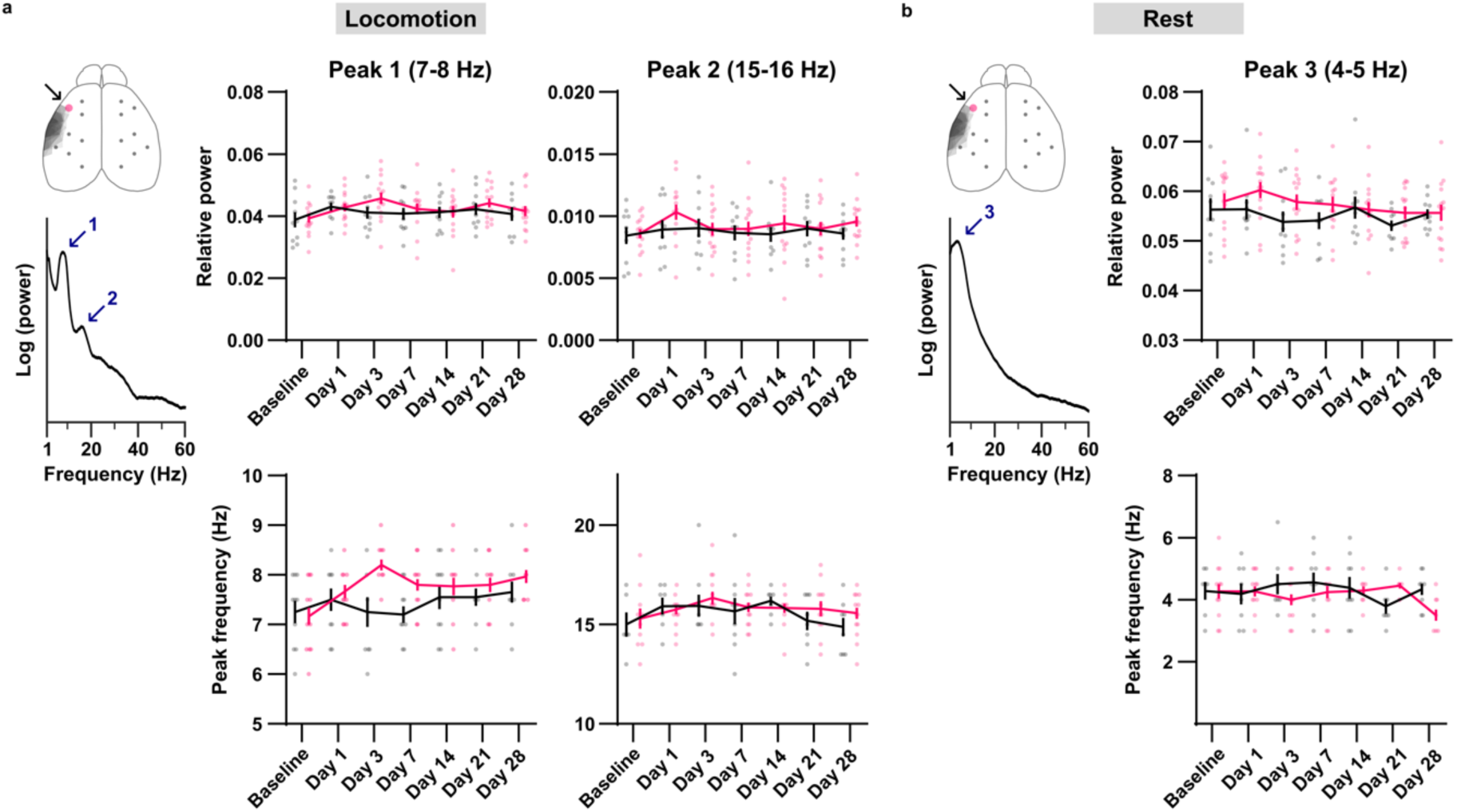
Relative power and peak frequencies of spectrum peaks in the motor cortex following pMCAO. **a**. Left, during locomotion two spectral peaks (blue arrows) were identified in M1 (black arrow, red dot). Middle, time course of relative power (top) and peak frequency (bottom) of the first spectral peak in stroke (red) and sham (black) animals. Right, time course of relative power (top) and peak frequency (bottom) of the second spectral peak in stroke (red) and sham (black) animals. Relative power was calculated through division of the absolute peak power by broadband power (1-60 Hz) of the spectrum. **b**. Same as a., shown for the resting condition. One spectral peak (blue arrow) was identified in M1 (black arrow, red dot). Stroke: n = 15, sham: n = 10. ** p < 0.01, * p < 0.05. Data is shown ± SEM.

## Supplementary tables

**Supplementary Tab. 1.** Structures affected by pMCAO stroke. Lesion hemisphere in brackets (L, left; R, right). MOp, primary motor area; SSp, primary somatosensory area; SSp-un, unassigned; SSp-m, mouth; SSp-ul, upper limb; SSp-n, nose; SSp-bfd, barrel field; SSs, supplemental somatosensory area; AUDd, dorsal auditory area.

| Mouse number<br>(hem.) | Anterior-posterior coordinates (in relation to bregma, mm) |  |  |  |  |  |  |  |
| --- | --- | --- | --- | --- | --- | --- | --- | --- |
|  | +1.5 | +1 | +0.5 | 0 | -0.5 | -1 | -1.5 | -2 |
| 1 (L) |  | SSp-m | SSp-m/n | SSs | SSs |  |  |  |
| 2 (R) | SSp-un | SSp-m | SSp-m/n<br>SSs | SSp-bfd<br>SSs | SSp-bfd<br>SSs | SSp-bfd<br>SSs | SSp-bfd<br>SSs | SSp-bfd |
| 5 (R) |  | SSp-m | SSp-m/n<br>SSs | SSp-bfd<br>SSs | SSp-bfd<br>SSs |  |  |  |
| 6 (L) |  | SSp-m | SSp-m/n<br>SSs | SSp-bfd<br>SSs | SSp-bfd<br>SSs | SSs | SSs |  |
| 11 (L) | SSp-un | SSp-ul/m | SSp-ul/m/n<br>SSs | SSp-bfd<br>SSs | SSp-bfd<br>SSs | SSp-bfd | SSs |  |
| 12 (R) |  | SSp-m | SSp-m/n<br>SSs | SSp-bfd<br>SSs | SSp-bfd<br>SSs | SSs | SSs |  |
| 13 (L) | SSp-un | SSp-m | SSp-m/n<br>SSs | SSp-bfd<br>SSs | SSs | SSs | SSs | AUDd |
| 14 (R) | SSp-un | SSp-ul/m | SSp-m/n<br>SSs | SSp-bfd<br>SSs | SSp-bfd<br>SSs | SSs | SSs |  |
| 18 (R) |  |  |  | SSs | SSp-bfd<br>SSs |  |  |  |
| 22 (R) |  | SSp-m | SSp-m/n<br>SSs | SSp-bfd<br>SSs | SSs |  |  |  |
| 27 (R) |  |  | SSs | SSp-bfd<br>SSs | SSp-bfd<br>SSs | SSp-bfd<br>SSs | SSp-bfd<br>SSs | SSp-bfd |
| 28 (R) |  |  | SSp-n<br>SSs | SSs | SSs |  |  |  |
| 29 (R) |  |  | SSp-n<br>SSs | SSp-bfd<br>SSs | SSp-bfd<br>SSs | SSs |  |  |
| 32 (R) |  | SSp-ul/m | SSp-ul/m/n | SSp-bfd<br>SSs | SSp-bfd<br>SSs | SSp-bfd<br>SSs | SSp-bfd<br>SSs |  |
| 33 (L) | SSp-un | SSp-ul/m | SSp-ul/m/n<br>SSs |  |  |  |  |  |

**Supplementary Tab. 2.** Correlation of infarct size with various parameters. Spearman r-(black, top) and p-values (grey, bottom) for correlation with SPR performance (% baseline), exponent and offset (relative to baseline), and broadband power (P1-60, power between 1 and 60 Hz, relative to baseline). n = 12. Loc, locomotion.

| Time<br>(day) |  | Spearman correlation of infarct size with: |  |  |  |  |  |  |
| --- | --- | --- | --- | --- | --- | --- | --- | --- |
|  |  | SPR<br>perf. | Exp.<br>(loc) | Exp.<br>(rest) | Offset<br>(loc) | Offset<br>(rest) | P1-60<br>(loc) | P1-60<br>(rest) |
| 1 | r | -0,41 | -0,27 | -0,10 | 0,17 | -0,02 | 0,06 | 0 |
|  | p | 0,19 | 0,40 | 0,77 | 0,59 | 0,96 | 0,87 | >0,99 |
| 3 | r | -0,58 | -0,21 | -0,27 | -0,03 | -0,05 | -0,52 | -0,22 |
|  | p | 0,05 | 0,51 | 0,40 | 0,94 | 0,89 | 0,09 | 0,50 |
| 7 | r | 0,11 | -0,20 | -0,20 | 0,03 | 0,04 | -0,20 | -0,18 |
|  | p | 0,73 | 0,54 | 0,53 | 0,94 | 0,90 | 0,53 | 0,57 |
| 14 | r | 0,24 | -0,08 | -0,10 | -0,21 | -0,17 | -0,25 | -0,17 |
|  | p | 0,46 | 0,80 | 0,75 | 0,51 | 0,60 | 0,44 | 0,60 |
| 21 | r | -0,23 | 0,03 | -0,08 | -0,51 | 0,01 | -0,48 | 0,05 |
|  | p | 0,47 | 0,92 | 0,80 | 0,09 | 0,99 | 0,12 | 0,89 |
| 28 | r | 0,11 | 0,14 | -0,08 | -0,31 | -0,17 | -0,42 | 0,02 |
|  | p | 0,73 | 0,67 | 0,81 | 0,32 | 0,60 | 0,18 | 0,96 |

**Supplementary Tab. 3.**
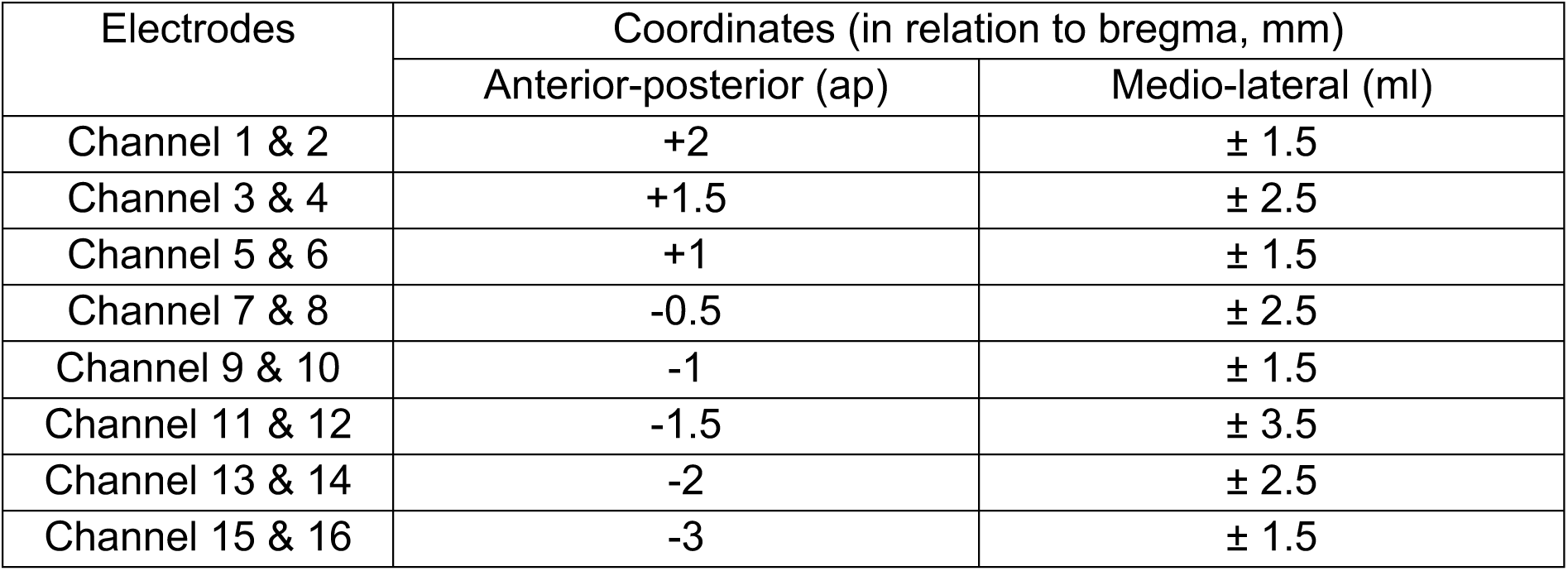
Stereotactic coordinates of implanted ECoG electrodes.

